# Identification of Regulatory Loci for Megakaryocyte and Hepatocyte Coagulation Factor V Expression in Mice

**DOI:** 10.1101/2025.08.14.670312

**Authors:** Adrianna M. Jurek, Kailey H. MacFadyen, Marisa A. Brake, Lacramioara Ivanciu, Rodney M. Camire, Randal J. Westrick

**Author notes:** Corresponding Author: Randal J. Westrick, 118 Library Dr., 305 Dodge Hall, Rochester, MI 48309.

## Abstract

**Background:** Factor V (FV) plays a central role in the coagulation cascade, acting in both a procoagulant and anticoagulant manner. The majority of FV is produced by the liver hepatocytes. In humans, FV is endocytosed by megakaryocytes, whereas in mice, FV is synthesized by megakaryocytes. Little is known about the genomic factors regulating FV transcription in humans and mice.

**Objective:** To investigate genomic regulatory mechanisms for coagulation FV levels in the hepatocytes and megakaryocytes of inbred mice.

**Methods:** Plasma and platelet FV levels were measured via ELISA in 5 mouse strains. A cross between the CAST/EiJ and DBA/2J strains was performed to generate 146 genetically informative F2 mice for analysis of circulating and platelet FV. Plasma and platelet FV levels were measured by ELISA for the F2 mice and whole genome genotyping for each F2 was performed using the TransnetYX MiniMUGA genotyping array. The genotyping and phenotyping data collected from these mice were then analyzed using quantitative trait loci (QTL) analysis.

**Results and Conclusions:** We identified one significant locus controlling plasma FV levels on Chromosome 1, ∼57.7 million base pairs upstream of the FV structural gene. We also identified a significant QTL for platelets on Chromosome 14 when sex was included as an interactive covariate, with additional suggestive loci present on Chromosomes 15 and 2. Our findings provide foundational information regarding the cell-type and sex-specific control of FV expression, establishing the basis for further investigations aimed at fine-mapping these loci and understanding how FV expression is regulated.

**Essentials:**

- Coagulation Factor V (FV, gene name *F5*) is primarily expressed in hepatocytes in humans but in hepatocytes and megakaryocyte/platelets in mice.
- Plasma and megakaryocyte/platelet *F5* expression varies significantly between inbred mouse strains.
- We identified significant loci controlling plasma and platelet *F5* expression in mice. Platelet *F5* expression is influenced by biological sex.

## Introduction

Factor V (FV) is a critical protein in the coagulation cascade. In its active form (FVa), it serves as a cofactor for activated factor X (FXa) in the prothrombinase complex, which produces thrombin and amplifies coagulation^1^. FV also has anticoagulant functions, acting with protein S (PS) as a cofactor for activated protein C (APC) to inactivate FVIIIa^2^, and with tissue factor pathway inhibitor (TFPI) to inhibit FXa^3^. In blood, FV is present in two compartments: 80% circulates in plasma and 20% is sequestered in platelet alpha-granules^4,5^. Human FV is produced by liver hepatocytes, contributing to the FV circulating in the plasma. Plasma FV is endocytosed by megakaryocytes and stored in platelet alpha granules^6,7^. In mice, hepatocytes produce circulating plasma FV^4^, and megakaryocytes produce platelet FV directly^8^. Platelet FV is thought to be more procoagulant than its plasma counterpart and increasing levels of platelet FV lead to decreased time to occlusion in a carotid artery injury mouse model^9^. Deficiency of FV causes mucosal lining and post-traumatic bleeding, and complete absence of FV is incompatible with life^10^.

Recently, *PLXDC2*, a cell surface transmembrane receptor, has been shown to correlate with the modulation of FV expression^11^ and *PSKH2*, a protein serine kinase, has been found to play a role in the variability of FV activity^12^, but otherwise little is known about the genomic regulation controlling expression of this critical protein. Here, we identified natural variations in the platelet and plasma FV levels of inbred mice and used QTL mapping to identify genomic loci regulating FV expression.

## Methods

### Mouse Strains

Five mouse strains were acquired from The Jackson Laboratory including: C57BL/6J (B6, Stock Number 000664), DBA/2J (DBA, Stock Number 000671), A/J (Stock Number 000646), CAST/EiJ (CAST, Stock Number 000928), and 129S1/SvImJ (129S1, Stock Number 002448). All mouse strains had an unmodified FV locus, are commonly used, and are phylogenetically distinct^13^.

### Generation of F2 Mice

Female DBA mice were outcrossed to male CAST mice to generate CASTD2F1 (F1) mice. Male and female F1 mice were intercrossed to generate the CASTD2F2 (F2) generation. The F2 mice were used in subsequent experiments. All mouse experiments followed Oakland University’s Institutional Committee on Use and Care of Animals guidelines.

### Isolation of Plasma

Whole blood was collected via cardiac puncture from B6 (n=15), DBA (n=26), A/J (n=17), 129S1 (n=14), CAST (n=12), F1 (n=19), and F2 (M=26, F=35) mice and diluted 9:1 in 3.2% buffered sodium citrate (Medicago, Aniara Diagnostica) using the techniques detailed in Brake et al^14^. Platelet poor plasma was stored at -80 °C until required for experiments.

### Isolation of Platelets

Platelets were isolated from B6 (n=18), DBA (n=14), A/J (n=12), 129S1 (n=17), CAST (n=13), F1 (n=20), and F2 (M=40, F=45) mice. Whole blood was diluted 20:1 with acid citrate dextrose/prostaglandin E1 (1mM cocktail) during exsanguination by cardiac puncture^14^. Blood was diluted and platelets were isolated as previously described in Siebert et al^13^.

### Genotyping by Mouse Universal Genotyping Array

Tail biopsies from all F2 mice, two DBA females, and two CAST males were submitted to TransnetYX for genotyping using their MiniMouse Universal Genotyping Array (MiniMUGA, Cordova, TN)^15^.

### Measurement of FV Levels in Plasma and Platelets

Total FV protein concentration was measured by enzyme-linked immunosorbent assay (ELISA) for plasma and platelets. Platelets were diluted 20x in RIPA buffer, and platelet protein concentration was determined by the Pierce BCA assay (ThermoFisher Scientific). Platelet protein concentration was normalized to 25 μg/mL using blocking buffer (1X PBS, 0.05% Tween-20, 6% BSA). Plasma was serially diluted with blocking buffer, 1:10 or 1:20 and then 1:25 for a final dilution of 1:250 or 1:500 depending on concentration. A Nunc Maxisorp (ThermoFisher Scientific) 96-well plate was coated with 50 μL of 1 μg/mL solution of GMA-755 rat anti-murine monoclonal FV antibody (Green Mountain Antibodies) in coating buffer (0.1M Carbonate pH 9.2) and incubated at 4°C overnight. The plate was washed 3 times with 200 μL/well wash buffer (0.01% Tween-20 in 1X PBS). 200 μL blocking buffer was incubated at room temperature for 2 hours, then washed. 50 μL of samples or varying amounts of recombinant murine FV (used for standard curve) purified from baby hamster kidney (BHK) cells^16,17^ were added in duplicate and incubated for 1 hour at 37 °C. The plate was then washed, and 100 μL of detection antibody (rat anti-murine FV GMA-752 antibody (100 μg/mL) labeled using HRP Conjugation Kit-Lightening Link (Abcam, diluted 1:4000 in blocking buffer)) was incubated at 37°C for 1 hour. The plate was then washed and 50 μL of 3,3′,5,5′-Tetramethylbenzidine (TMB) was added, and the plate was incubated in the dark for 10 minutes at room temperature. The reaction was quenched with 25 μL 1N H_2_SO_4_. The plate was read immediately using the Biotek Synergy H1 plate reader at 450 nm and standards were plotted on a log scale. Final measurements for platelet FV were calculated to ng/μg of platelet protein, and plasma FV values were multiplied by the dilution factor.

### Quantitative Trait Loci Analysis

Separate genome scans of the platelet and plasma F2 mice were performed using the R/qtl package^18^ using the methods from Siebert et al^13^ and Hailey-Knott regression to determine if platelet and plasma FV regulatory loci exist. There were 3,523 and 3,538 informative markers used for the plasma and platelet QTL analysis, respectively. Initial analysis was performed using the GRCm38 mouse build, and chromosome positions were converted to the GRCm39 build using UCSC LiftOver (https://genome.ucsc.edu/cgi-bin/hgLiftOver).

### Statistical Analyses

The graphed results are shown as mean ± SEM. Graphs were generated using GraphPad Prism, and statistical analysis was performed using ANOVA adjusted for multiple comparisons for groups 3 or larger, and Welch’s t-test was used for groups of two. A p-value of <0.05 was considered significant.

## Results and Discussion

### FV Antigen Levels

Analysis of plasma FV (Figure 1A) revealed significant differences between several of our assayed strains. Although CAST (132.6 μg/mL) and DBA (119.2 μg/mL) strains had similar intrastrain plasma FV levels when both sexes were grouped together, we observed significant sex-specific differences in FV antigen levels between the female CAST and DBA mice (p=0.0044, Figure 1B).

**Figure 1:**
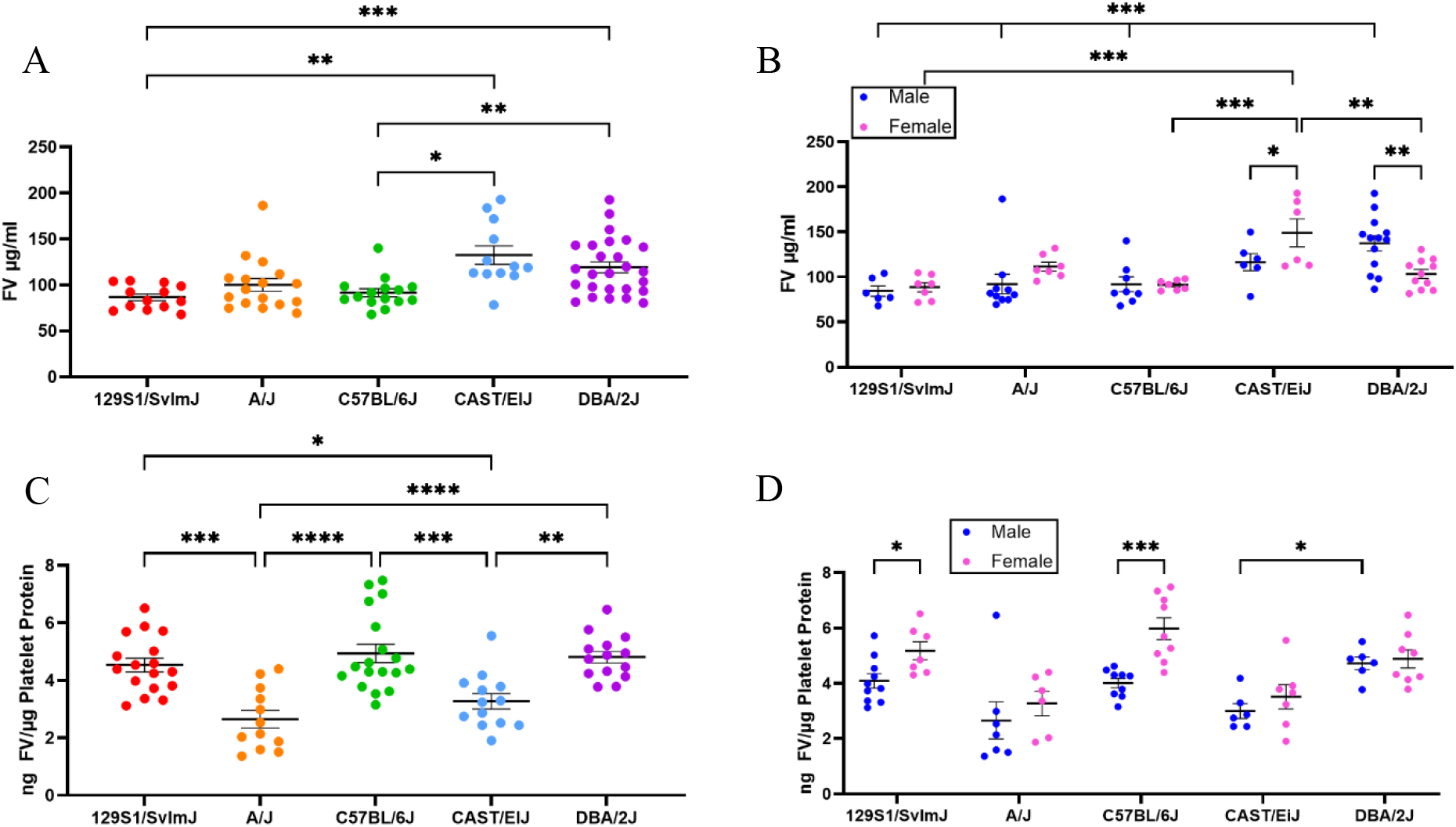
Factor V Levels in Plasma (A, B) and Platelets (C, D) from Five Inbred Mouse Strains. FV protein levels were measured via ELISA in platelet-poor plasma and washed platelet lysates from 129S1, A/J, B6, CAST, and DBA strains. A) Plasma FV levels were significantly different between 129S1 and CAST (p=0.0065), B6 and CAST (p=0.0168), B6 and DBA (p=0.0059) and 129S1 and DBA (p=0.0004). B) Plasma FV levels were significantly different between the males and females in the DBA (p=0.0011) and CAST (p=0.0236) strains. There was also a significant difference between the DBA females and the CAST females (p=0.0044), DBA and 129S1, A/J and B6 males (p=0.0004, 0.0003, and 0.0009 respectively), and CAST vs 129S1 and B6 females (p=0.0003 and 0.0006 respectively). C) Platelet FV levels were significantly different between 129S1 and A/J (p=0.0001), 129S1 and CAST (p=0.0166), A/J and B6 (p<0.0001), A/J and DBA (p<0.0001), B6 and CAST (p=0.0005), and CAST and DBA (p=0.0033). D) Platelet FV levels in females were significantly higher within the 129S1 (p=0.0322) and the B6 (p=0.0001) strains. We also observed a significant difference between the male CAST and male DBA mice (p=0.0336). Data are presented as mean ± SEM. Statistical comparisons were performed using Brown-Forsythe and Welch ANOVA for plasma FV (A), Ordinary 1-way ANOVA for platelet FV (C), and 2-way ANOVA for sex differences (B, D) with multiple comparisons adjusted for with Dunnett’s T3, Tukey’s and Šídák’s post hoc tests as appropriate.

Platelet FV antigen levels were highly variable between the five strains (Figure 1C), with the DBA and CAST strains having mean FV levels of 4.814 and 3.277 ng/μg platelet protein, respectively (p=0.0033). A significant sex difference was also seen between the male CAST and DBA mice (p=0.0336, Figure 1D). Prothrombin time (PT) and activated partial thromboplastin time (aPTT) were not significantly different between any strains, despite the significant differences in plasma FV antigen levels^19^.

There were no significant differences between the F1s and F2s in the plasma or platelets, or between the males and females in the plasma (Figure 2A-C). Interestingly, though there was no sex difference in the parental platelet FV levels, a sex difference appeared in the F2 generation (p<0.0001, Figure 2D).

**Figure 2:**
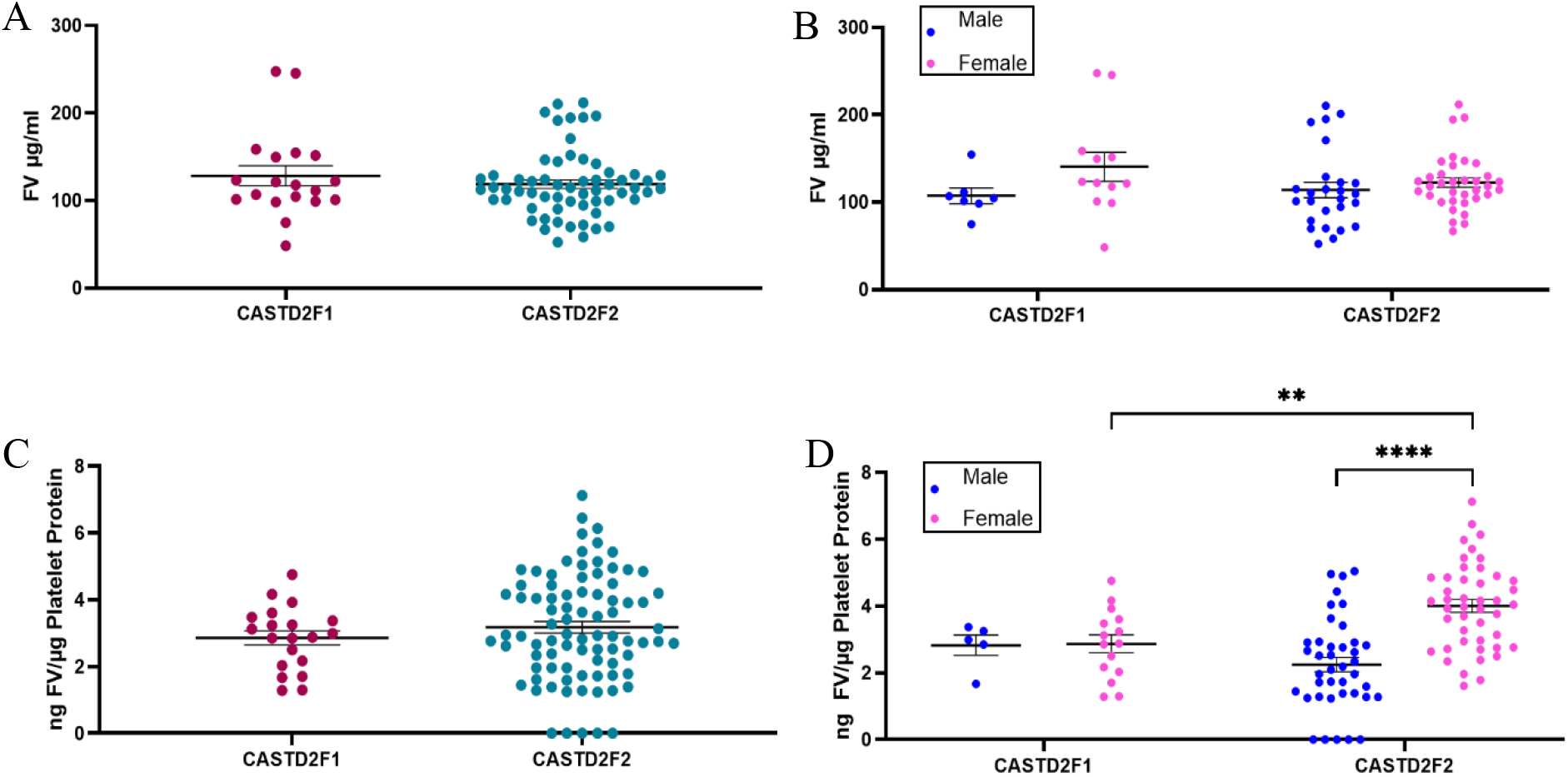
FV Antigen Levels in F1 and F2s in the Plasma and Platelets. FV protein levels were measured by ELISA in platelet-poor plasma and washed platelet lysates from F1 and F2 mice. A) There was no significant difference between the F1 and F2 strains. B) There was no significant sex difference within the strains. C) There was no significant difference between the strains. D) There was a significant difference in platelet FV antigen levels between the F2 males and females (p<0.0001) and between the F1 females and the F2 females (p=0.0177). Data are presented as mean ± SEM. Statistical comparisons were performed using Welch’s Corrected two-tailed t test (A), Unpaired two-tailed t test (C), and 2-way ANOVA (B, D).

### Identification of a Significant Chromosome 1 Regulatory Locus for Plasma FV

To identify genomic regions regulating plasma FV levels, QTL analysis was performed on 61 F2 mice. A significant peak was observed on Chromosome (Chr) 1 (LOD 5.24, significance threshold 4.21, p=0.00641, Figure 3A (Black graph)) from 95,095,725-166,889,809. At the maximum peak on Chr 1 (main peak: 151,294,711 marker gUNC1944082, range: 144,425,338-153,204,726), mice homozygous and heterozygous for the DBA allele were associated with lower plasma FV when compared to those homozygous for the CAST allele (p<0.0001 and p=0.0006, respectively, Figure 3B). When considering sex as an additive and interactive covariate in a full model (Figure 3A (Blue Graph)), an increase was observed in the LOD score at Chr 1 position 106,272,710 (max range: 106,272,710-116,168,330, LOD 6.79, significance threshold 6.34, p=0.0246, Figure 3C). At the maximum peak in the full model, mice homozygous and heterozygous for the DBA allele were associated with significantly lower FV levels than those homozygous for the CAST allele (p=0.0001, data not shown), similar to the initial QTL analysis results. Additionally, we observed a significant difference between the male and female mice heterozygous for CAST/DBA at our full model peak (p=0.0416, Figure 3D).

**Figure 3:**
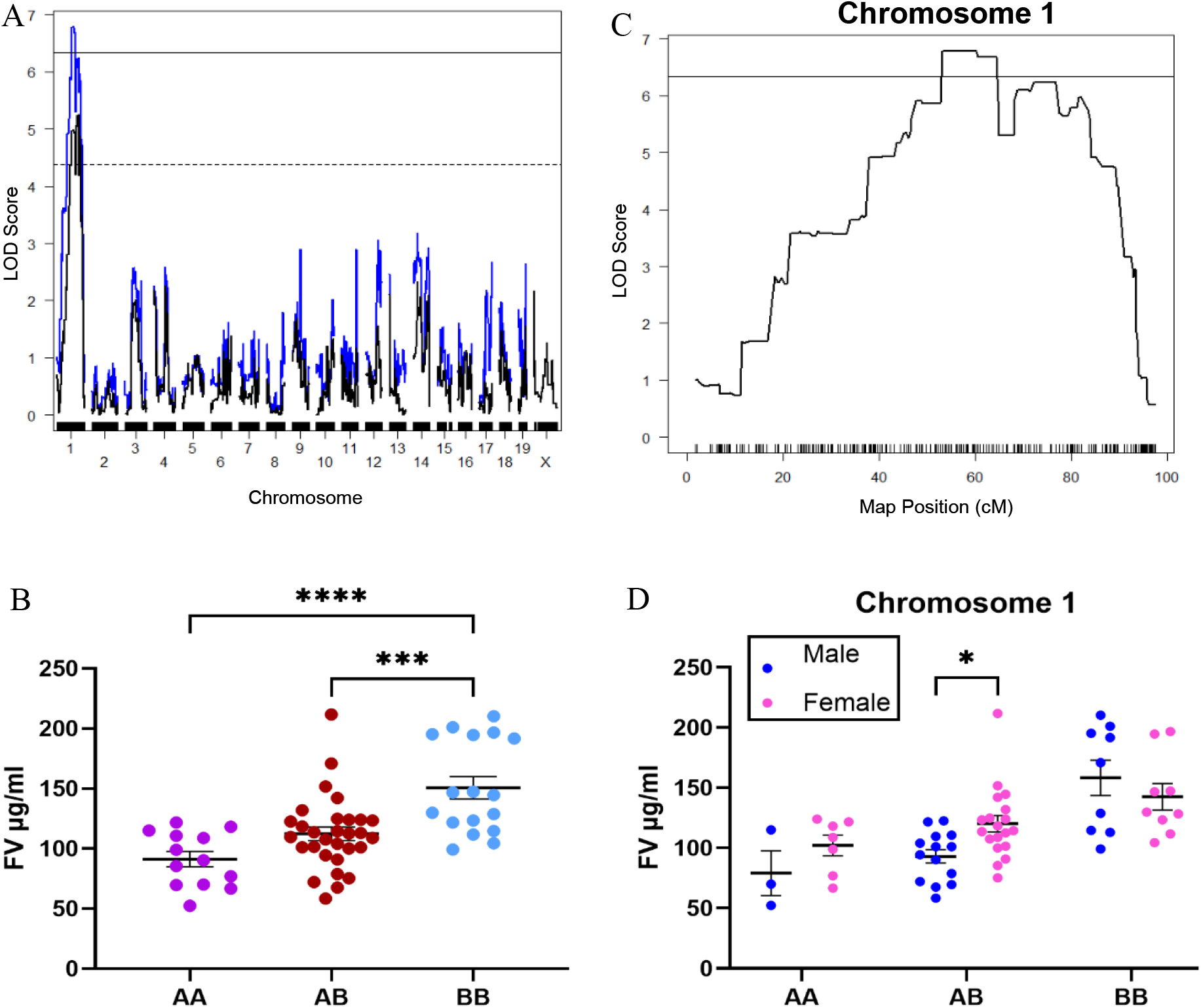
Plasma CASTD2F2 QTL Results. QTL results were obtained from running *scanone* and Hailey-Knott regression on r/qtl to identify loci responsible for controlling plasma FV. Sex was used as an additive and interactive covariate. Solid line indicated significance threshold for full model, dotted line indicates significance threshold for initial model. A) Initial QTL (black) and full model QTL (blue) with a significant peak identified on Chr 1 with a LOD score 5.24, significance threshold 4.21, p=0.00641 (initial) and LOD 6.79, significance threshold 6.34, p=0.0246 (Full Model). No suggestive peaks were seen. B) Zoomed in view of Chr 1 peak from full model with significance threshold. C) Effect plot demonstrating that individuals homozygous for CAST (BB) at marker gUNC1944082 had significantly higher plasma FV when compared to those homozygous for DBA (AA, p<0.0001) and heterozygous (AB, p=0.0006). D) In the full model, we saw a significant difference between the males and females in the mice heterozygous for CAST/DBA (p=0.0416) at the most significant peak. Data are presented as mean ± SEM. Statistical comparisons were performed using Ordinary 1-way ANOVA (C) and 2-way ANOVA (D) adjusted for multiple comparison using Tukey’s and Šídák’s post hoc tests as appropriate.

Interestingly, the initial QTL identified the significant peak at Chr 1 without including sex as a covariate, despite our findings of sex specificity in the parentals. This could be due to the female CAST vs DBA strain difference seen in Figure 1B. When sex as a covariate was included, the peak strengthened, which was expected given our initial findings.

The mouse FV gene is located on Chr1:163,979,407-164,047,846 (MGI, GRCm39), which is included in the minimal significant recombinant interval of our QTL model. However, the main significance peak is in a transcriptionally rich region 57,706,697 bp upstream of the gene. Within that region is a serpin cluster, which includes *Serpinb2* (PAI-2), a known component of the coagulation cascade^20,21^. While there is no direct linkage with FV, it is possible that alteration of PAI-2 levels impacts the overall fibrinolytic balance with indirect effects on FV. However, due to the large number of potential causative variants located in the significant range, additional studies are required to identify the precise genetic regulator.

### Identification of Sex Specific Platelet Regulatory Loci on Chromosomes 14 and 15

To identify genomic regions regulating platelet FV levels, QTL analysis was performed on 85 F2 mice. While no significant or suggestive peaks were detected with the initial QTL scan, after using sex as an interactive covariate (Figure 4A), a significant peak was observed on Chr 14 at position 21,755,370-23,755,370 (LOD 3.45, significance threshold 2.32, p=0.00641, Figure 4B), with the most significant peak located at position 23,079,378. A suggestive peak was observed on Chr 15 at position 4,758,870-7,086,683 (LOD 1.64, suggestive 20% threshold 1.58, p=0.17844, Figure 4C), with the highest peak marker located at 4,758,870. At the peak on Chr 14, male mice homozygous for DBA and CAST had lower platelet FV levels than females (p<0.0001, Figure 4D). At Chr 15 peak, the same trend was seen, as well as heterozygous females also having significantly higher FV than males (heterozygous: p=0.0068, homozygous DBA: p<0.0001, homozygous CAST: p=0.0013, Figure 4E).

**Figure 4:**
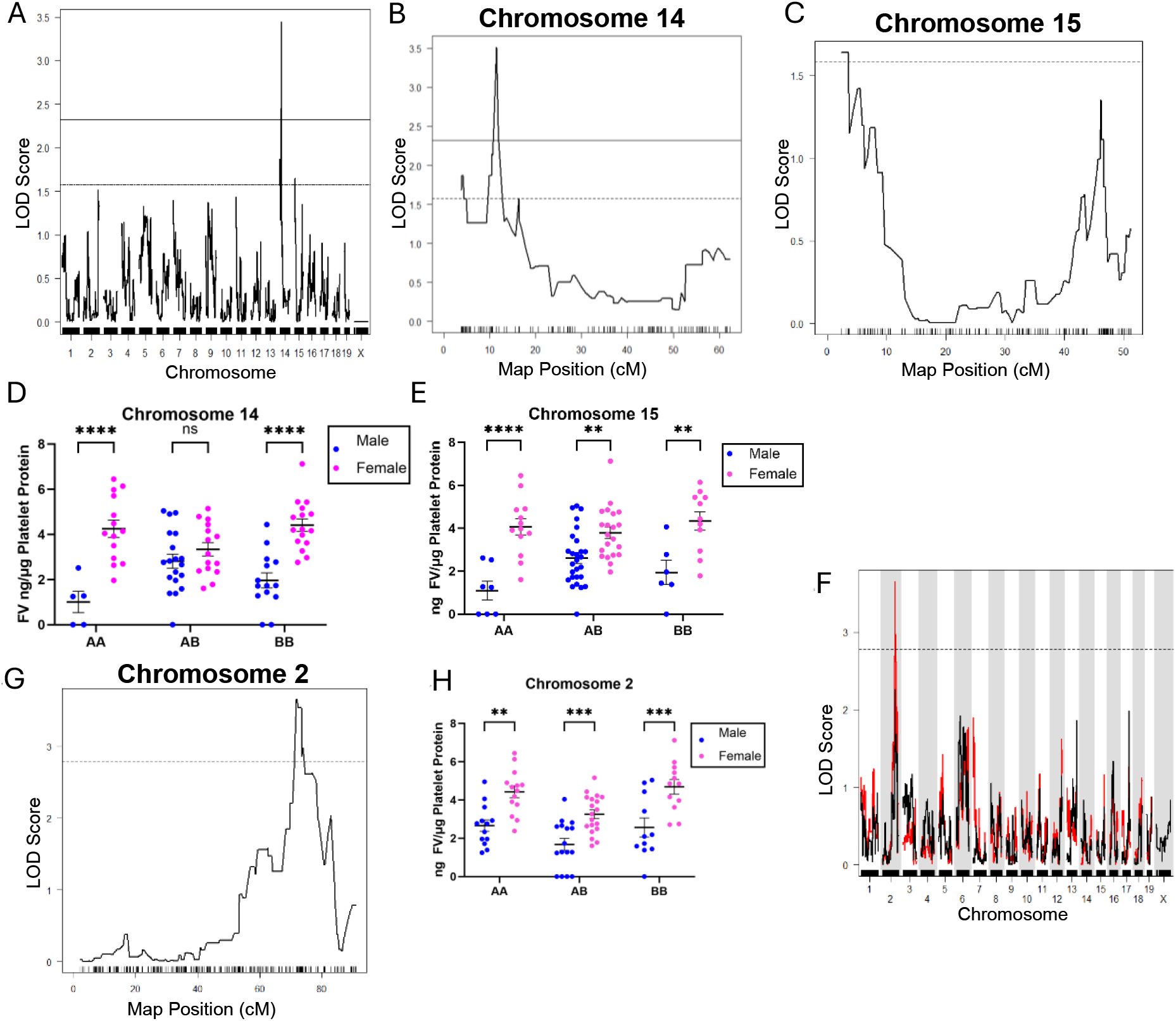
Platelet CASTD2F2 Whole QTL with Sex as an Interactive Covariate (A), and QTL with Sex as an Additive Covariate (F). QTL results were obtained from running *scanone* and Hailey-Knott regression on R/qtl to identify loci responsible for controlling platelet FV. Sex was used as an additive and interactive covariate. Solid line indicated significance threshold, dotted line indicates suggestive threshold. A) Whole platelet QTL with sex as an interactive covariate. A significant peak was seen on Chr 14, and a suggestive peak was found on Chr 15. B) Magnified Chr 14 peak, LOD score 3.42, significant threshold 2.32, p=0.00641. C) Magnified Chr 15 peak, LOD Score 1.64, suggestive 20% threshold 1.58, p value= 0.17844. D) Individuals homozygous for DBA (AA) and CAST (BB) at marker gUNC23694515 on Chr 14 showed a significant difference in sex (p<0.0001). E) The same difference was seen at Chr 15 marker gUNC24956990 (AA p<0.0001, BB p=0.0013) but the significant sex difference was seen in heterozygous individuals as well (AB p=0.0068). F) Whole QTL with sex as an additive covariate with a suggestive peak on Chr 2. G) Magnified Chr 2 peak, LOD 3.66, suggestive threshold 2.78, p=0.113. H) At marker gUNC4139799 on Chr 2, the same trend was seen, with males having statistically lower FV levels than the females at each genotype (AA p=0.0011, AB p=0.0009, and BB p=0.0003). Data are presented as mean ± SEM. Statistical comparisons were performed using 2-way ANOVA (D, E, and H) and multiple comparisons were adjusted for using Šídák’s post hoc test.

When sex was used as an additive covariate (Figure 4F), a suggestive Chr 2 peak at position 143,502,220-146,651,920 was present (LOD 3.66, suggestive 50% threshold 2.78, p=0.113, Figure 4G). The highest LOD score in the region was marker gUNC4139799, located at 143,901,800. Consistent with the interactive covariate scan, male mice had FV levels significantly lower than females for all genotypes (homozygous DBA: p=0.0011, heterozygous: p=0.0009, homozygous CAST: p=0.0003, Figure 4H). While the FV interacting coagulation gene *TFPI*^22^ (Chr2:84,263,199-84,307,119, MGI, GRCm39), and *PLXDC*2 (Chr2:16,361,115-16,760,650, MGI, GRCm39) are both located on Chr 2, they are a significant distance from our mapped region, suggesting that any potential effects of these loci would be due to higher order interactions of the type precipitated by the spatial organization of chromatin^23^. Since these platelet regulatory loci were identified when investigating sex as a covariate due to the significant sex difference seen in our F2s, a potential role of sex hormones, like estrogen, in platelet FV level variation may be present.

## Conclusions

Our study is the first to demonstrate strain-dependent differences in FV expression among laboratory mice. We identified a significant locus regulating plasma FV levels on Chr 1 in a region containing a serpin cluster. We also identified one significant and two suggestive loci on Chrs 14,15, and 2 respectively, that may play a role in regulating platelet FV levels in a sex specific manner.

Recently, Munsch et al. used a multi-omics approach to identify genetic regulators of plasma FV levels in humans. They identified several candidate genes and proteins, 4 of which fall proximally to the loci identified here^24^. The *PLXDC2* (Chr 2), *TFPI* (Chr 2), *COX6C* (Chr 15), and *SURF1* (Chr2)^24^ genes/proteins were associated with FV levels. While outside of our minimal significant regions in this initial mapping experiment, they could still possibly regulate FV levels in mice.

Future work aimed at fine mapping and identifying the regulatory elements controlling platelet and plasma-specific FV expression will increase our understanding of cell-type specific gene regulation and may reveal novel targets or strategies for modulating FV in humans.

